# Testing relationships between multimodal modes of brain structural variation and age, sex and polygenic scores for neuroticism in children and adolescents

**DOI:** 10.1101/2019.12.20.883959

**Authors:** Linn B. Norbom, Jaroslav Rokicki, Dennis van der Meer, Dag Alnæs, Nhat Trung Doan, Torgeir Moberget, Tobias Kaufmann, Ole A. Andreassen, Lars T. Westlye, Christian K. Tamnes

## Abstract

Human brain development involves spatially and temporally heterogeneous changes, detectable across a wide range of magnetic resonance imaging (MRI) measures. Investigating the interplay between multimodal MRI and polygenic scores (PGS) for personality traits associated with mental disorders in youth may provide new knowledge about typical and atypical neurodevelopment. We derived independent components across cortical thickness, cortical surface area, and gray/white matter contrast (GWC) (n=2596, 3-23 years), and tested for associations between these components and age, sex and-, in a subsample (n=878), PGS for neuroticism. Age was negatively associated with a single-modality component reflecting higher global GWC, and additionally with components capturing common variance between global thickness and GWC, and several multimodal regional patterns. Sex differences were found for components primarily capturing global and regional surface area (boys>girls), but also regional cortical thickness. For PGS for neuroticism, we found weak and bidirectional associations with a component reflecting right prefrontal surface area. These results indicate that multimodal fusion is sensitive to age and sex differences in brain structure in youth, but only weakly to polygenic load for neuroticism.

## Introduction

The cerebral cortex and subjacent white matter undergo substantial modifications and refinement during childhood and adolescence ^1-3^, paralleled by the qualitative and quantitative evolution of cognitive abilities ^4, 5^. These developmental changes coincide with increasing risk for mental disorders ^6, 7^ Mapping variation in brain development using magnetic resonance imaging (MRI) may therefore not only inform ontogenetic models of neurocognitive development, but also delineate aberrant spatiotemporal patterns related to emerging psychopathology.

Cortical development is multifaceted, with cortical volume and its subcomponents thickness and surface area showing distinct developmental patterns ^8-10^, and genetic underpinnings ^11, 12^ Less explored measures of signal intensity variation in the Tl-weighted image may reflect additional and partly distinct neurobiological properties ^13-16^. Specifically, the contrast between cortical grey matter (GM) and closely subjacent white matter (WM) intensities, the grey/white matter contrast (GWC), has been shown to be heritable ^16^, and associated with development ^14^, aging ^13, 17^, and schizophrenia ^18^. Although the underlying biology of GWC is complex and debated, the measure has been linked to differential intracortical and WM myelination.

The onset of puberty, as well as the pubertal period, differs between adolescent boys and girls ^19, 20^. There have also been reports of sex specific shifts in the onset of developmental psychopathology, as well as variations in overall risk for mental disorders ^7, 21^. Results concerning both the extent, and metric specificity of sex related neurodevelopmental differences are however, inconclusive ^8, 9, 22^ and no studies have investigated sex related developmental differences in GWC.

The relationships between distinct brain structural properties captured by different MRI measures are poorly understood, but might be informed by methods such as linked independent component analysis (LICA), which can parse the common and unique variance across multiple modalities into separate components of shared variance ^23^. LICA studies have reported unique structural patterns sensitive to lifespan development ^24, 25^, and psychopathology ^26, 27^ Multimodal fusion, also including GWC, thus shows promise for capturing both typical brain developmental patterns and patterns associated with susceptibility for mental illness.

High levels of neuroticism is associated with several forms of psychopathology ^28, 29^, including the overarching “p factor” from dimensional psychopathology models ^30, 31^, and internalizing symptoms from core domain models ^28^, which captures vulnerability toward mood and anxiety disorders ^31, 32^ Twin and family studies have shown that genetic differences account for about 40% of the trait variance ^33^, and genome-wide association studies (GWAS) have documented a highly polygenic signal ^34^, in line with most complex traits ^35^. Polygenic scores (PGS) are defined as the weighted sum of an ensemble of trait-associated alleles ^36^, and reflect the individual level of cumulative genetic signal across the genome. Only one prior neuroimaging study has investigated the association between PGS for neuroticism and brain structure, reporting negative associations with regional cortical surface area in two adult samples. Studies in youth are lacking, but may provide insight into the brain structural correlates of genetic dispositions for broad psychopathology-associated traits in a period of life when many mental disorders typically emerge.

To this end, we combined cortical thickness, surface area, and GWC using FLICA (FMRIB’s LICA) in 2596 youths (3-23 years), and tested for associations between the resulting multimodal modes of variation and age, sex and-, in a subsample (n=878), PGS for neuroticism and two of its subcomponents (depressed affect and worry). Based on an extant literature on child and adolescent brain development ^9, 14, 38, 39^ we hypothesized that age would be particularly associated with components dominated by GWC and thickness. Next, we predicted associations between sex and modes of variation reflecting surface area ^40, 41^. Finally, based on findings from a previous study ^37^ we hypothesized that the strongest associations between PGS for neuroticism, depressed affect and worry, would be with modes reflecting surface area.

## Materials and Methods

### Participants

We combined data from two large-scale publicly available US samples; the Pediatric Imaging, Neurocognition, and Genetics (PING) study (http://ping.chd.ucsd.edu) and the Philadelphia Neurodevelopmental Cohort (PNC) (Permission No. 8642). PING is a multisite initiative, consisting of typically developing youths aged 3-21 years with genotyping, standardized behavioral measures, and multimodal imaging for a large subgroup (n = 1239) ^4, 42, 43^. PNC is a population-based sample consisting of 8-23 year old youths with genetic, cognitive and clinical data, as well as multimodal imaging for a large subgroup (n=1601) ^44, 45^. The cohorts, including recruitment and exclusion criteria, are further described in Supplementary Materials.

We first excluded PNC subjects with severe medical health conditions (n=70), before combining all subjects with available imaging data from both samples (n=2727). After stringent MRI quality control, 115 participants were excluded due to missing or poor quality T1 data (see below), and 16 additional subjects were removed due to missing demographics or scan site information. The final imaging sample thus consisted of 2596 subjects (1316 girls) aged 3.0-23.2 years (mean=13.8, SD=4.6) (Figure 1). See Supplementary Table 1 for demographics.

**Figure 1.**
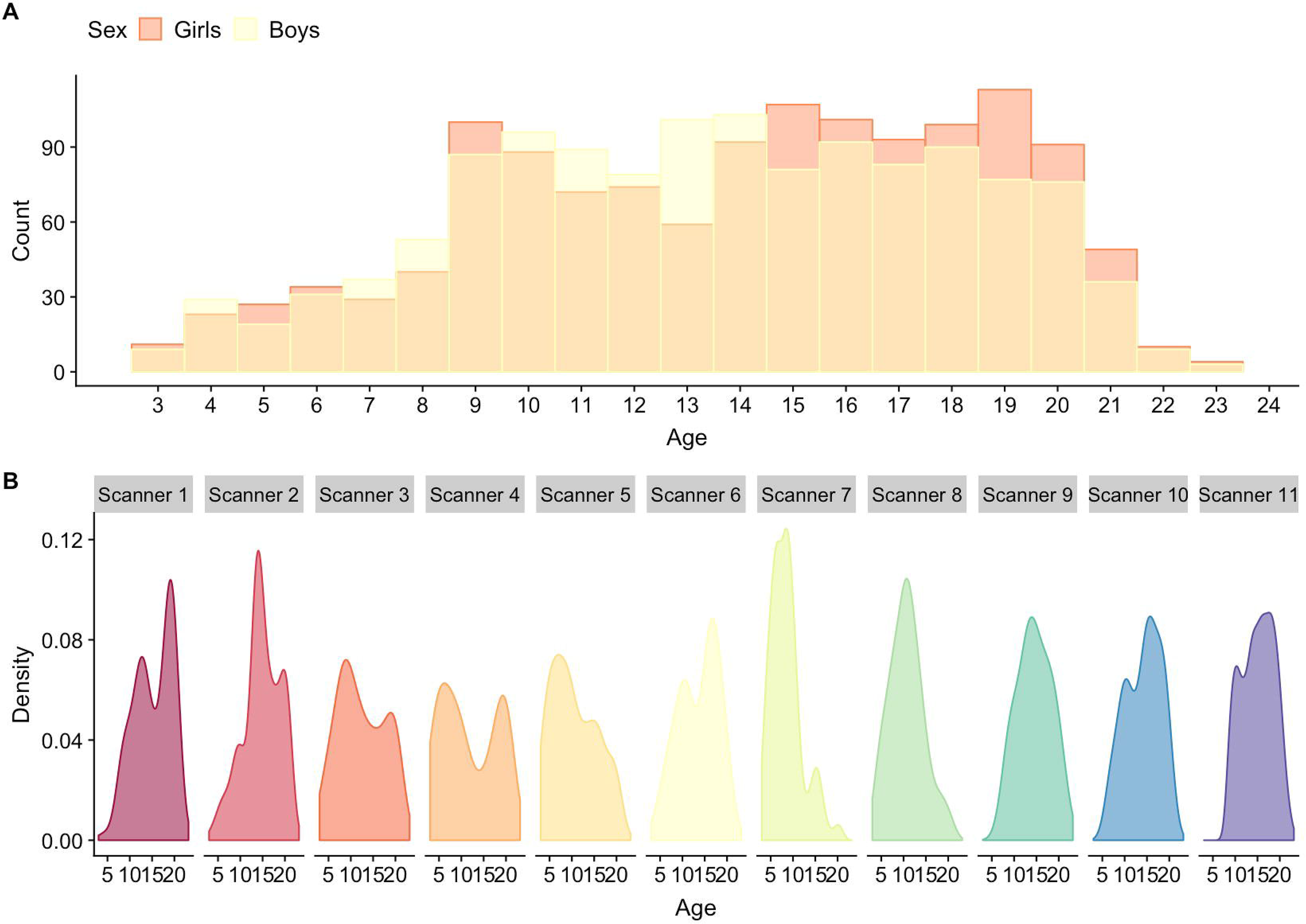
Age and sex distribution for the full MRI sample (n=2596). (A) Depicts the age and sex distribution for the full MRI sample, while (B) depicts the age distribution within each scanner. Scanner 1-10 are scanners employed by the PING study, while scanner 11 was used in the PNC.

### MRI acquisition and processing

The included imaging data was acquired on 11 separate 3T whole-body scanners from Philips Medical Systems (Achieva), GE medical systems (Signa HDx and Discovery MR750), and Siemens (TrioTim), the latter including a single scanner on which the full PNC sample was scanned. T1 voxel sizes ranged from approximately 0.9 to 1.2 mm. See Supplementary Materials for detailed descriptions of acquisition parameters, and care and safety procedures implemented for scanning children.

The T1-weighted datasets were processed using FreeSurfer (FS) 5.3 (http://surfer.nmr.mgh.harvard.edu), which performs volumetric segmentations and cortical surface reconstructions, including the “white”-(grey/white matter boundary), and “pial”-(grey/ cerebrospinal fluid (CSF) boundary) surface ^46, 47^ Cortical thickness was computed as the shortest vertex-wise distance between the white and pial surface, while cortical surface area, based on the white surface, was computed by the amount of vertex-wise expansion and contraction needed to fit a common template (fsaverage) ^46, 47^ As implemented previously ^14, 18, 48^ and fully described in Supplementary Materials, we extracted intra-subject signal intensities from the non-uniform intensity normalized volume (nu.mgz). GWC was then computed as: 100 x (white - gray)/[(white - gray)/2] ^18^, so that higher GWC reflects greater GM/WM signal intensity difference. Thickness, area and GWC surface maps were registered to fsaverage and smoothed using a Gaussian kernel of 15-, 10-, and 10-mm full width at half maximum (FWHM), respectively, to obtain similar smoothness across modality surfaces ^26^

See Supplementary Materials and previous descriptions ^14, 49^ for in detail accounts of the stringent quality evaluation performed on the imaging data to confirm consistency across sites and as in-scanner head motion is common in children.

### FLICA

Thickness, area, and GWC surface maps were decomposed into spatially independent modes of variation using FLICA ^23, 25^ (http://fsl.fmrib.ox.ac.uk/fsl/fslwiki/FLICA). As imaging data had been obtained from various scanners, and initial investigations showed large scanner effects, we residualized all surface maps for effects of scanner before running FLICA. This was done by use of generalized additive models, using the “gam” function in R (https://www.r-project.org/), with 10 knots and age and sex as additional covariates. Age, sex and model residuals were returned to surface maps. See Supplementary Figures 1-3 and Supplementary Table 2 for comparisons of surface maps before and after scanner residualization. FLICA was thereafter run with 3000 iterations, and a final model order of 60, which was pragmatically chosen based on the highest cophenetic coefficient ^50^, as compared with model orders of 40 and 80. See Supplementary Figure 4 for an ordered bar plot of the explained variance of each component.

### Polygenic scores

We calculated PGS in a subset of participants with white European ancestry (n= 878), using the PRSice v1.25 software, based on an earlier GWAS on neuroticism ^51^. PGS p-thresholds were calculated within the range of 0.001 and 0.5, using default settings, including removal of the major histocompatibility complex (MHC; chromosome 6, 26-33Mb) and pruning of SNPs based on linkage disequilibrium and p-values. We first chose a significance threshold of 0.05 based on the convention of several previous GWAS studies ^34^, and additionally, based on a previous implementation ^52^, performed a principal component analysis (PCA) in R (https://www.r-project.org/) on all p-thresholds. The first factor explained 88.1% of variance, with the p-threshold 0.329 showing the highest contribution, and was extracted as a complementary and considerably more liberal PGS p-threshold.

Based on results from the GWAS ^51^ reporting that neuroticism’s genetic signal partly originated in two distinguishable sub-clusters termed “depressed affect” and “worry”, we additionally calculated PGS for these using the same approach as described above. In addition to the standard 0.05 threshold, the first PCA factor explained 93.4% of the variance for both, with p-thresholds 0.257 and 0.253 showing the highest contribution for depressed affect and worry, respectively.

### Statistical analyses

We used linear models, to test for main effects of age, sex and PGS for neuroticism on all FLICA components in R. First, for each of the 60 components we tested for linear effects of age, with scanner as an additional co-variate. Then we tested for main effects of sex, covarying for scanner and age. Finally, we tested for main effects of the different polygenic scores on the FLICA components. Effects were tested on each of the 60 components in separate models, with scanner, age, sex, and the first 4 PCA derived genetic principal components included in the model to account for population stratification. P-values from all models within each test were adjusted for multiple comparisons by false discovery rate using Hochberg’s procedure and a significance threshold of 0.05.

To visualize each model, we residualized covariates from component loadings by linear regression, and plotted them using ggplot2 ^53^. See Supplementary Materials for ANOVA analyses, performed to investigate possible effects of scanner.

In order to allow for comparisons with more conventional unimodal approaches, we examined the vertex-wise effects of age, sex and the PGS for neuroticism sum score on each of our modalities separately, using GLM as implemented in the Permutation Analysis of Linear Models (PALM) toolbox ^54^ (see Supplementary Materials).

## Results

### Multimodal components

Among the 60 independent components (ICs), IC1 explained 34.3% of the variance in the included modalities across the 2596 subjects, and was dominated by global GWC (97%) (Figure 2). The spatial pattern showed relatively lower weighting in insula, sensorimotor and visual cortices and higher weighting within prefrontal cortices. IC2, which explained 14.8% of variance, was dominated by global surface area (94%), with higher weighting within prefrontal, occipital, and lateral temporal regions (Figure 3). IC3, explained 11.7% of variance, and captured global cortical thickness (55%) and global negative GWC (45%). The cortical thickness spatial pattern showed higher weighting in frontal regions, pre-and postcentral sulcus, and within certain parietal and occipital folds. The GWC pattern showed higher negative weighting in insula, medial visual- and parietal regions (Figure 2).

**Figure 2.**
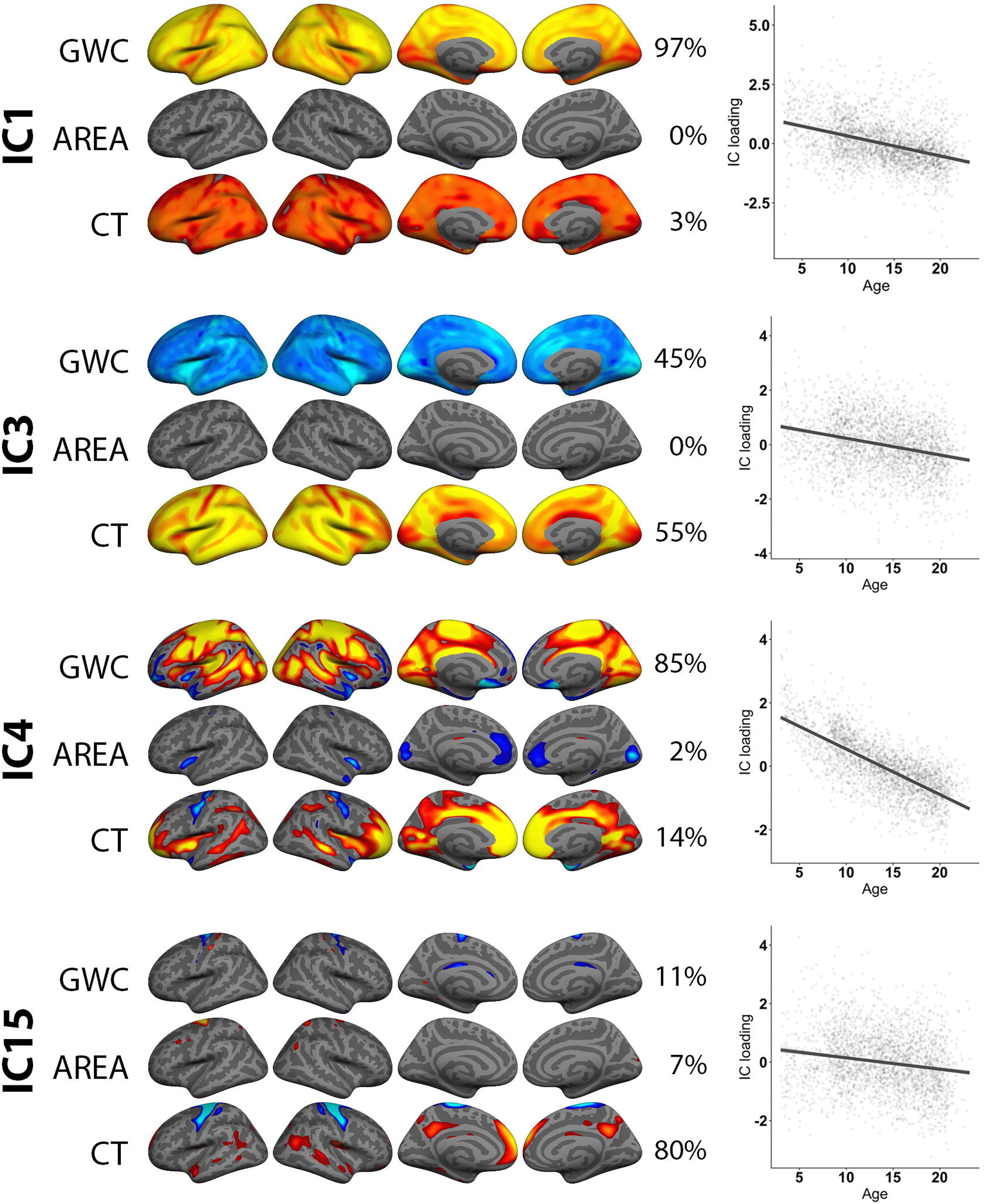
Associations between age and multimodal imaging independent components (ICs). The figure depicts FMRIB’s linked independent component analysis (FLICA) weighted spatial maps for the four ICs with the strongest age associations. The accompanying plots show linear models of age plotted against scanner residualized IC loading. In general, all components were thresholded within a minimum and maximum of 8 and 17 standard deviations (SD) respectively, except for the global maps within IC1, IC3 and IC4, which were thresholded with a higher maximum SD value in order to reveal nuances in the global pattern.

**Figure 3.**
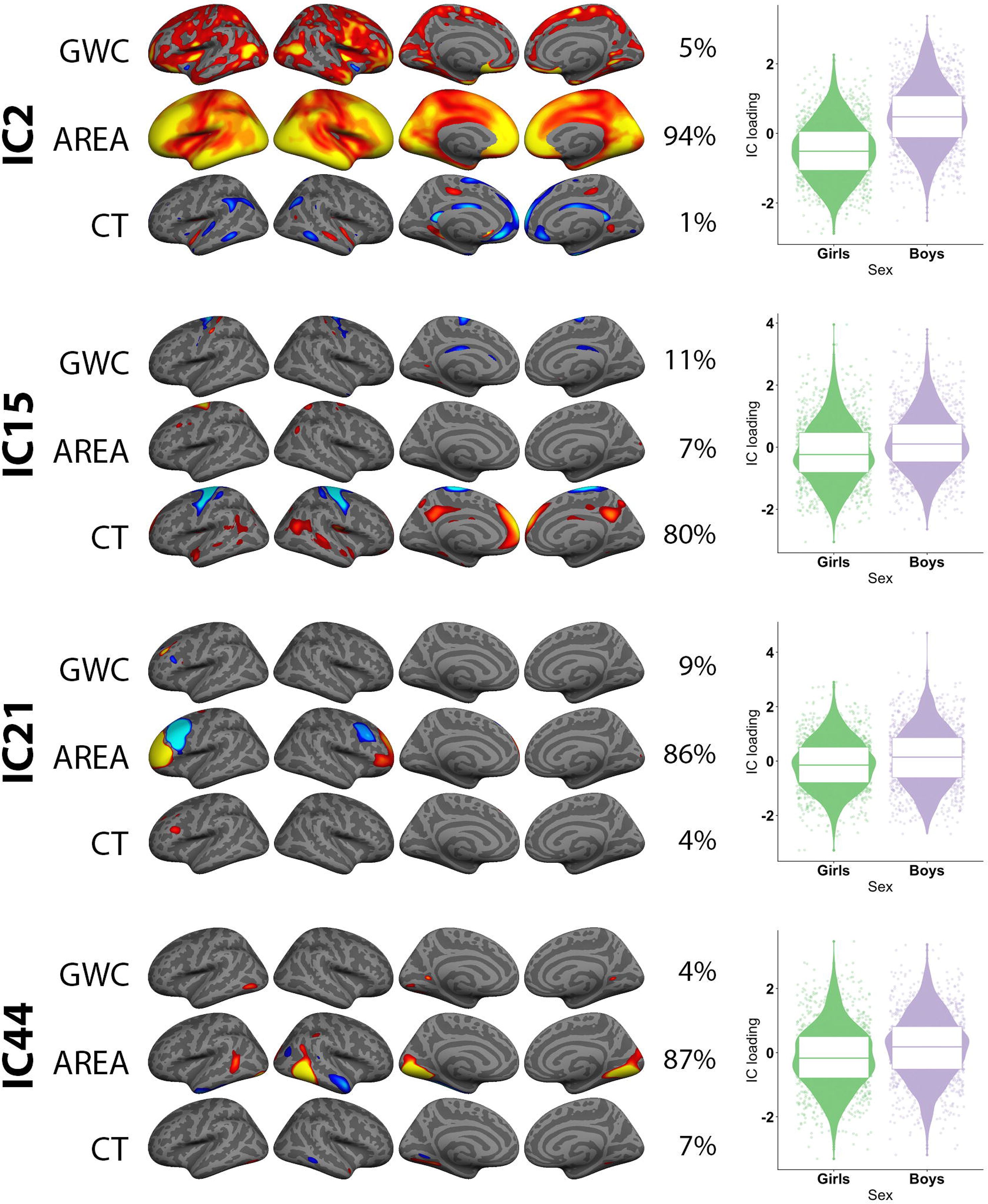
Associations between sex and multimodal imaging independent components (ICs). The figure depicts FMRIB’s linked independent component analysis (FLICA) weighted spatial maps for the four ICs with the strongest effect of sex. The accompanying violin boxplots shows sex plotted against scanner and age residualized IC loading. All components were thresholded within a minimum and maximum of 8 and 17 standard deviations (SD) respectively, except for the global map within IC2, which was thresholded with a higher maximum SD value in order to reveal nuances in the global pattern.

IC4 explained 6.0% of the variance, while the remaining 56 components in sum explained 33.2%. IC4-IC60 were single-modality or multimodal, regionally specific, and often bilateral. Multimodal components largely captured corresponding regions across modalities, and in these instances (with the exception of IC3), GWC and thickness generally showed directional weighting correspondence, which was opposite to surface area (i.e. higher thickness and GWC with smaller surface area).

ANOVA on age residualized component loadings revealed that none of the components were highly sensitive to scanner differences (Supplementary Table 3), with IC4 showing the only significant scanner effect (F= 4.3, adjusted p = <0.001).

### Associations between multimodal components and age

Linear models revealed significant age associations for 17 of the 60 components when including scanner in the model (Table 1, Supplementary Figure 5 and 6). The four components with the strongest age effects are shown in Figure 2. With higher age, IC1 revealed globally lower GWC, IC3 globally lower thickness coupled with globally higher GWC, and IC4 bilaterally lower GWC across large regions of the brain including cingulate, insula, occipital, and pre- and postcentral cortices extending into frontal regions, and higher GWC in minor rostral middle frontal and temporal pole regions. IC15 revealed bilaterally thicker precentral cortices, as well as thinner medial prefrontal and posterior cingulate cortices with higher age.

**Table 1.** Overview of age and sex effects on multimodal imaging components. The table shows the T statistics for age (controlling for scanner) and sex (controlling for scanner and age) effects on each independent component, as well as P-values adjusted for multiple comparisons by false discovery rate correction. P-values at or below 0.05 are marked in bold.

| IC | Age<br>effect<br>(t) | P-value | Sex<br>effect<br>(t) | P-value |
| --- | --- | --- | --- | --- |
| 1 | -24.86 | <.001 | 0.08 | 0.939 |
| 2 | -3.68 | **0.011** | 29.25 | <.001 |
| 3 | -16.88 | <.001 | -3.39 | **0.026** |
| 4 | -69.34 | <.001 | -2.34 | 0.612 |
| 5 | -8.09 | <.001 | 1.44 | 0.939 |
| 6 | -5.32 | <.001 | 0.53 | 0.939 |
| 7 | -6.22 | <.001 | 7.33 | <.001 |
| 8 | -2.06 | 0.924 | -5.25 | <.001 |
| 9 | 1.29 | 0.924 | 4.15 | **0.002** |
| 10 | 3.74 | **0.009** | 1.76 | 0.939 |
| 11 | 5.44 | <.001 | -0.29 | 0.939 |
| 12 | 4.86 | <.001 | 3.62 | **0.012** |
| 13 | -8.08 | <.001 | -2.38 | 0.574 |
| 14 | -3.79 | **0.007** | -3.12 | 0.066 |
| 15 | -10.18 | <.001 | 8.47 | <.001 |
| 16 | 2.04 | 0.924 | -4.25 | **0.001** |
| 17 | 0.23 | 0.924 | 0.98 | 0.939 |
| 18 | -0.54 | 0.924 | -4.94 | <.001 |
| 19 | -0.92 | 0.924 | -0.19 | 0.939 |
| 20 | 0.77 | 0.924 | -3.54 | **0.016** |
| 21 | -1.63 | 0.924 | 7.36 | <.001 |
| 22 | -3.15 | 0.068 | 1.95 | 0.939 |
| 23 | -0.62 | 0.924 | -0.8 | 0.939 |
| 24 | -0.87 | 0.924 | -0.39 | 0.939 |
| 25 | -1.42 | 0.924 | -0.34 | 0.939 |
| 26 | 1.06 | 0.924 | -4.27 | **0.001** |
| 27 | 1.76 | 0.924 | 2.26 | 0.732 |
| 28 | -0.3 | 0.924 | -1.57 | 0.939 |
| 29 | 3.15 | 0.068 | 3.95 | **0.004** |
| 30 | 0.68 | 0.924 | -0.92 | 0.939 |
| 31 | -3.59 | **0.015** | 0.39 | 0.939 |
| 32 | -0.09 | 0.924 | 0.87 | 0.939 |
| 33 | -0.33 | 0.924 | -1.08 | 0.939 |
| 34 | 2.52 | 0.453 | 4.21 | **0.001** |
| 35 | 0.3 | 0.924 | 3.78 | **0.007** |
| 36 | 2.67 | 0.295 | -1.16 | 0.939 |
| 37 | -0.11 | 0.924 | 3.03 | 0.086 |
| 38 | -0.4 | 0.924 | 2.08 | 0.939 |
| 39 | -0.33 | 0.924 | 5.34 | <.001 |
| 40 | -0.72 | 0.924 | -1.61 | 0.939 |
| 41 | 5.18 | <.001 | -0.74 | 0.939 |
| 42 | 1.73 | 0.924 | -0.56 | 0.939 |
| 43 | 3.04 | 0.096 | 5.97 | <.001 |
| 44 | 3.27 | **0.047** | 7.63 | <.001 |
| 45 | 0.86 | 0.924 | 2.65 | 0.279 |
| 46 | -1.95 | 0.924 | 3.83 | **0.006** |
| 47 | -3.21 | 0.057 | -3.9 | **0.004** |
| **48** | -3.91 | **0.005** | 0.59 | 0.939 |
| **49** | -1.9 | 0.924 | -0.72 | 0.939 |
| **50** | -1.74 | 0.924 | 3.53 | **0.016** |
| **51** | 1.88 | 0.924 | -3.7 | **0.009** |
| **52** | 1.31 | 0.924 | -0.51 | 0.939 |
| **53** | -0.48 | 0.924 | -0.92 | 0.939 |
| **54** | -1.43 | 0.924 | 2.2 | 0.826 |
| **55** | 0.44 | 0.924 | 1.45 | 0.939 |
| **56** | -0.22 | 0.924 | -1.86 | 0.939 |
| **57** | 2.02 | 0.924 | -1.6 | 0.939 |
| **58** | -0.73 | 0.924 | 1.9 | 0.939 |
| **59** | 2.13 | 0.924 | 3.49 | **0.019** |
| **60** | -0.37 | 0.924 | -6.34 | **<.001** |

The multimodal results highly concurred with results from unimodal analyses (see Supplementary Materials and Supplementary Figure 7).

### Associations between multimodal components and sex

Linear models revealed significant sex associations for 24 components when including age and scanner in the models (Table 1, Supplementary Figure 5 and 6). The four components with the strongest sex effects are shown in Figure 3. For boys as compared to girls IC2 revealed globally larger surface area, with strongest effects within prefrontal, occipital and lateral temporal cortices, and IC15 bilaterally thinner precentral cortices, as well as thicker medial prefrontal and posterior cingulate cortices. Moreover, for boys as compared to girls IC21 revealed larger lateral rostral prefrontal surface area, and smaller caudal prefrontal surface area, and IC44 larger surface area within a bilateral, mainly medial occipital region.

The multimodal results highly concurred with results from unimodal analyses (see Supplementary Materials and Supplementary Figure 8).

### Associations between multimodal components and PGS for neuroticism

In the white European ancestry sub-group (see Supplementary Table 1), linear models including scanner, age, sex, and 4 genetic principal components representing population stratification revealed no significant associations with any of the PCA-based PGS. The p<0.05 SNP selection threshold revealed a significant association between PGS for neuroticism with IC23 (t= −4.2, adjusted p= 0.004), which seemed to be driven by the PGS for depressed affect (t= −3.7, adjusted p= 0.016) (Figure 4). IC23 reflects a pattern with larger surface area within a right hemisphere ventrolateral prefrontal region and smaller surface area within an adjacent superior prefrontal region. There was no associations between the PGS worry scores thresholded at 0.05 and any of the multimodal components.

**Figure 4.**
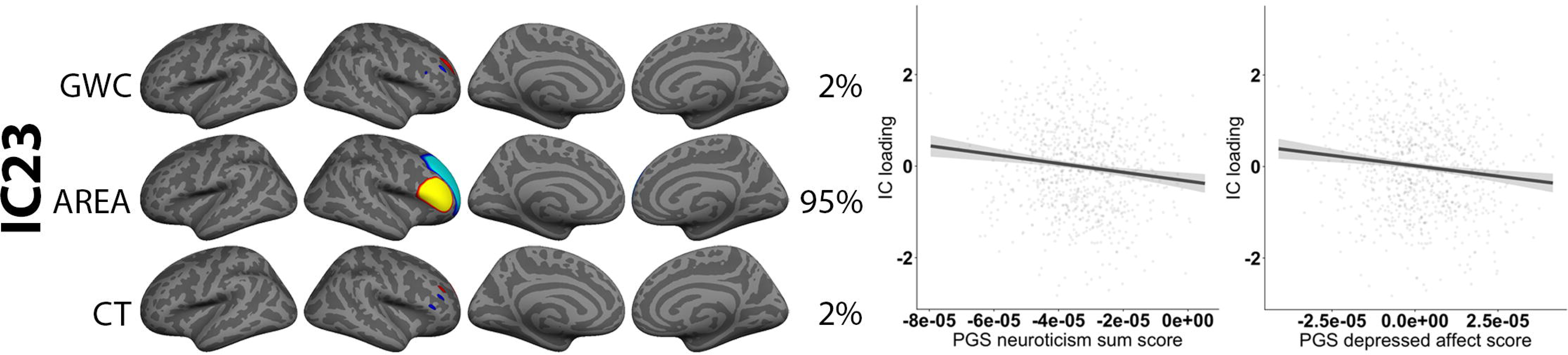
Association between polygenic scores (PGS) for neuroticism and a single multimodal imaging independent component (IC). The figure depicts FMRIB’s linked independent component analysis (FLICA) weighted spatial maps for the IC with an effect of PGS. The accompanying plots show linear models of PGS score plotted against scanner, age-, sex, and population stratification-residualized IC loading. The IC was thresholded within a minimum and maximum of 8 and 17 standard deviations (SD) respectively.

There were no significant unimodal associations between PGS for neuroticism and any of the imaging modalities. This was as expected, as frontal areal associations from LICA were modest, and vertex-wise analyses require stringent correction for multiple testing.

## Discussion

This multimodal structural brain imaging study of 2596 participants aged 3-23 years yielded three main results. First, higher age was most strongly associated with lower global GWC, with some regional variation. Higher age was also associated with globally thinner cortex, in particular encompassing prefrontal regions. Second, several components showed sex differences; boys had larger global and regional surface area compared to girls, and some regional cortical thickness differences. Third, there was a significant association between polygenic scores for neuroticism and a component encompassing surface area in prefrontal regions in the right hemisphere. Comparisons to more conventional vertex-wise unimodal approaches showed high correspondence, while additionally demonstrating unique benefits of LICA as it modeled relationships between several unimodal patterns.

The imaging component explaining the largest amount of variance was global and dominated by GWC, indicating a prominent role of GWC in childhood and adolescent neurodevelopment. Several of the other early components also had substantial contribution from GWC. Moreover, regions overlapping across modalities showed directional weighting correspondence between GWC and thickness, which was the opposing direction to area. As intracortical myelin is predominantly found in deeper cortical layers^55^, and our GM sampling strategy was a percentage, sampling of thicker cortex may to a larger degree include voxels in more superficial cortical layers. This could also result in a larger contrast between GM and WM. Our results are also in accordance with prior reports of negative associations between cortical thickness and surface area change^9, 56^.

Several imaging components were sensitive to age and, as hypothesized, the strongest age associations were seen for components dominated by GWC and thickness. Converging with prior studies^13, 14^, we found a negative association between age and a global GWC component. This indicates that GM and WM intensities appear more similar with higher age. Although a decidedly indirect measure, there is evidence supporting that the lowering of GWC is partly due to brightening of cortex, which concurs with other cortical intensity studies^15, 49^, possibly through the developmental process of intracortical myelination, which continues across and beyond adolescence^57^. Moreover, a component capturing global cortical thickness was also negatively associated with age, in line with previous studies^8, 9, 40, 43, 58^. Intriguingly, the negative association between cortical thickness and age was co-modeled with globally relatively *higher* GWC. This could be an addition to previous findings showing that the apparent cortical thinning in development is due to an underestimation of the true cortical thickness. The rationale is that axonal myelination mostly occur within deep cortical layers where encroaching white matter penetrates the cortical neuropil^58, 59^ As myelin brightens the appearance of the cortex, deep myelinated cortical layers could be misclassified as WM, thereby shifting the grey/white boundary outward^58, 60, 61^. Indeed, a cross-sectional study investigating magnetization transfer and cortical thickness in youths aged 14-24 reported that association regions were thicker and less myelinated compared to primary sensory regions in early adolescence, but that during development these regions had faster rates of thinning and myelination^62^. Moreover, a recent study investigating GM and WM changes within visual cortex using quantitative MRI and mean diffusivity, reported that tissue growth, and specifically increased levels of myelin as verified by postmortem cortical data, underlay the apparent cortical thinning in development^59^.

As hypothesized, sex differences were most pronounced for components encompassing cortical surface area, and as previously reported, boys showed overall larger surface area as compared to girls^8^. Beyond this scaling effect, boys had increases in regional occipital and prefrontal surface area, as well as bilaterally thinner precentral cortices and thicker medial prefrontal and posterior cingulate cortices, as compared to girls. Prior studies on cortical sex differences have been divergent^63^, but stable differences in surface area across age is a consistent finding. The present study indicates that there are few sex differences in GWC in youth.

Exploratory analyses on the associations between PGS for neuroticism and multimodal modes of brain structural variation yielded limited findings. The results showed a bidirectional association between PGS for neuroticism, as well as PGS for depressed affect, and an imaging component capturing prefrontal surface area within the right hemisphere. One prior study has investigated the association between PGS for neuroticism and brain structure in adults, using a discovery (n ≈ 1000) and replication sample (n≈ 600). In the discovery sample, PGS for neuroticism was associated with decreased regional surface area, including within inferior frontal gyrus, which overlaps with the region reported in the current paper, but this result was not replicated in the discovery sample^37^. Other studies have found associations between phenotypic neuroticism and smaller frontal surface area^64, 65^, and in youth a positive association between emotional stability and surface area, albeit in temporal lobe regions^66^. However, a recent large-scale study (n ≈ 1000) attempting to detangle the inconsistent findings within neuroimaging personality research, reported no significant associations between any of the Big Five personality traits and any of the structural measures, including cortical surface area or thickness ^67^

The present study attempted to investigate the genetic disposition of- and not phenotypic neuroticism, and it should be stated that PGS have been reported to explain little of the known heritability in complex traits. PGS based on the GWAS used in the current paper have been reported to explain a maximum of 4.2% of the variance in neuroticism^34^. Similarly, a combined meta-analysis GWAS of neuroticism using almost 10,000 participants from the UK Biobank cohort, as well as 2 separate replication cohorts, reported that PGS derived from the UK Biobank sample only captured about 1% of the variance in neuroticism in the replication cohorts. In addition, most of the genome-wide significant alleles were not independently replicated ^68^. Furthermore, an attempt to apply PGS to split-half samples failed to significantly predict any of the tested personality traits of the study, including neuroticism ^69^ Beyond descriptions of-, and comparisons to several prior findings, we therefore refrain from making strong interpretations of our PGS results.

There are several limitations to the current study. First, reported GWC associations cannot be directly credited to myelination in general or intracortical myelination specifically. Although cholesterol in myelin is a major determinant of T1-weighted signal intensity^70, 71^, so in part is iron, water content and dendrite density^72-74^. While cortical GM intensity corresponds closely with histologically based myelin profiles^75^, GWC is microstructurally complex and a product of GM and WM. Intricate combinations of both matter types can therefore produce changes in GWC. Second, the cross-sectional design is suboptimal for the study of development^76^.

To conclude, our results indicated that in childhood and adolescence, GWC explains more variance in brain structure across subjects than cortical thickness or surface area. Further, negative age-related associations were found for global GWC and cortical thickness, the latter in combination with regionally relatively higher GWC. These results are in accordance with biological interpretations of intracortical myelination and an outward shifting of the grey/white border with increasing age. Boys had larger global and regional cortical surface area than girls, and there were also regional sex differences in cortical thickness. Finally, we found weak bidirectional associations between PGS for neuroticism and regional prefrontal surface area in the right hemisphere. Future developmental longitudinal studies are needed to validate the consistency of the reported effects.

## Acknowledgments and disclosures

The current work was supported by the Research Council of Norway (223273, 249795, 230345, 288083), the South-Eastern Norway Regional Health Authority (2014097, 2016083, 2019069), and the European Commission’s 7th Framework Programme (602450, IMAGEMEND). The Pediatric Imaging, Neurocognition and Genetics Study (PING) in part provided data. The National Institutes of Health (Grant RC2DA029475), and the National Institute on Drug Abuse and the Eunice Kennedy Shriver National Institute of Child Health & Human Development funded collection and sharing. PING data are disseminated by the PING Coordinating Center at the Center for Human Development, University of California, San Diego. Data was also in part from the Philadelphia Neurodevelopment Cohort (PNC), and support for this collection was provided by grant RC2MH089983 to Raquel Gur, and RC2MH089924 to Hakon Hakonarson. The participants were recruited through the Center for Applied Genomics at The Children’s Hospital in Philadelphia.

