## Supplemental materials for "Testing relationships between multimodal modes of brain structural variation and age, sex and polygenic scores for neuroticism in children and adolescents"

**Supplementary Materials and Methods**

*Participants*

Concerning PING, subjects within the desired age range and with fluent English abilities were recruited across the US, through local postings and outreach facilities. Exclusion criteria included somatic illness, preterm birth, diagnosis of mental disorders (not including ADHD), mental retardation, and contraindications for MRI ^1^. For PNC, participants were recruited through a previous separate Children’s Hospital of Philadelphia-study enrollment ^2^. Exclusion criteria included, mental retardation, non-English proficiency, medical problems that could affect brain function or neuroimaging tasks, claustrophobia, and contraindications for MRI ^3^.

### Care and safety procedures during image acquisition

The PING sample included children as young as 3 years old, and it should be stated that no subjects were sedated for imaging^4^. Subjects from the PING sample were accompaniment by a parent or a technician into the scanning environment, and exposed and habituated to the scanner before image acquisition. There were also opportunities to rest in between scans, or other behavioral support if needed. During scanning, all subjects were asked to lay as still as possible and the youngest subjects were given extra head padding. During the structural parts of the protocol, participants were presented with a movie of their choice with sound delivered via headphones^4^. Before image acquisition, subjects from the PNC sample were presented with a mock scanning session with a retired MRI scanner and head coil. These scannings were accompanied with auditory recordings of the noise produced by gradient coils for each pulse sequence. The mock scan was performed in an attempt to familiarize the youths with the MRI environment, and to practice laying very still^3^.

*MRI acquisition*

To account for challenges associated with multisite imaging of children, an optimized standard imaging protocol was implemented for the PING study, across vendors and models to obtain similar contrast properties and image-derived quantitative measures^1^. Acquisition protocols with pulse sequence parameters identical to or nearly identical to those implemented at UC San Diego were installed on each scanner. At UC San Diego, data were obtained on a GE 3T Signa HDx scanner and a 3T Discovery 750x scanner (GE Healthcare, Waukesha, WI) using eight-channel phased array head receiver coils. The protocol included a sagittal 3D inversion recovery spoiled gradient echo (IR-SPGR) T1-weighted volume optimized for maximum gray/white matter contrast with prospective motion correction (PROMO). TE = 3.5 ms, TR = 8.1 ms, TI = 640 ms, flip angle = 8°, receiver bandwidth = ± 31.25 kHz, FOV = 24 cm, freq = 256, phase = 192, slice thickness = 1.2 mm **^5^**. Scan durations for the T1 sequence was 8 minutes and 5 seconds. Voxel sizes ranged from ≈ 0.9-1.2mm^3^.

For PNC, signal excitation and reception was obtained using a quadrature body coil for transmit and a 32-channel head receiver coil. Gradient performance was 45mT/m, with a maximum slew rate of 200 T/m/s. T1 weighted imaging was obtained using a magnetization prepared rapid acquisition gradient-echo (MPRAGE) sequence (TR=1810 ms; TE=3.51 ms; FoV=180×240 mm; Resolution=0.94×0.94×1.0 mm). Receive coil (i.e. B1) shading was reduced by selecting the Siemens pre-scan normalize option, which corrects for B1 inhomogeneity based on a body coil reference scan. Total scan time of the entire protocol was 50 minutes and 32 seconds. All scans were acquired with a straight magnet axial orientation (i.e. non-oblique)^3^.

*MRI quality assessment*

Both PING and PNC imaging data were assessed by the same individuals and with automated flagging procedures, followed by visual inspection.

For PING, a few subjects were first directly excluded due to having unsatisfactory T1 resolution: > 1.2mm voxel size in any direction (n=2). A few subjects were also directly excluded as they had multiple runs within the same folder, highly suspected of being different individuals (n=4). All NIFTI images including several duplicate images, unique re-runs from the same session, and images from the same subject but at different sessions i.e. after re-positioning in the scanner, were then processed through the quality assessment pipeline MRIQC^6^. This program returns flagging of poor images as well as a numeric quality index based on measures of noise, spatial and tissue distribution, artifacts such as motion, and sharpness. After visual inspection, flagged subjects were either included, or excluded due to poor image quality (n=46) when there was no option to replace the image with another satisfactory run. For non-flagged subjects with several images, the highest quality sequence was chosen based on having superior “quality index” number which is outputted from MRIQC.

For PNC, to assess the quality of Freesurfer cortical reconstructions, we implemented a flagging procedure based on robust principal component analysis for detecting signal-to-noise and segmentation outliers^7, 8^. Flagged datasets were carefully inspected and minor edits were performed when necessary (n = 459) before re-running cortical reconstructions. Subjects with poor and corrupt image quality were thereafter excluded (n=63).

*GWC calculation*

We extracted intra-subject signal intensities from the non-uniform intensity normalized volume (nu.mgz) using the mri_vol2surf function. Vertex-wise GM intensities were sampled at six equally spaced points, starting 10% from the white surface and extending a maximum of 60% into the cortical ribbon. We selected this endpoint to minimize partial volume effects from voxels containing CSF. Vertex-wise WM intensities were sampled at 10 equally spaced points, starting 0.15 mm below the white surface and extending the fixed distance of 1.5 mm into subcortical WM. To obtain single vertex-wise intensity values for GM and WM separately, we averaged the intensity values from all sampling points. GWC was then computed as: 100 x (white - gray)/[(white - gray)/2] ^9^

*Statistical analyses*

Before main analyses, we investigated possible components sensitive to scanner effects, by the use of ANOVA as implemented in R (<https://www.r-project.org/>) on FLICA loadings, which we residualized for age by linear regression. All p-values were adjusted for multiple comparisons by false discovery rate using Hochberg’s procedure, and a significance threshold of 0.05.

In order to allow for comparisons with more conventional unimodal we used general linear models as implemented in the Permutation Analysis of Linear Models (PALM) toolbox ^10^. We first tested the linear effect of age on each of our modalities separately, i.e. vertex-wise GWC, cortical surface area and cortical thickness, with scanner included as a co-variate. We then tested for the effects of sex on our unimodal measures, covarying for scanner and age. Finally, in a subgroup of subjects with white European ancestry (n=878), we tested the effects of the PGS for neuroticism sum score thresholded at 0.05, on all modalities separately, with scanner, age, sex, and the first four principal component analysis (PCA) derived components included in the model to account for population stratification. To assess statistical significance we used 10,000 permutations and family wise error (FWE) correction and a significance threshold of p<0.05, after correcting for both hemispheres.

**Supplementary Results**

*Associations between unimodal vertex-wise structure and age*

Permutation testing revealed a strong and near global negative association between age (Supplementary Figure 7) and vertex-wise GWC, indicating lower contrast across most of the cortex with higher age. The differential pattern in association strength was highly similar to IC1 as well as IC4. Insular regions showed a positive association with age, which could correspond with IC4 findings. Linear models revealed moderate associations between age and vertex-wise surface area, indicating smaller area with higher age in several and mostly anterior cortical regions, as well as larger surface area within insula and precentral gyrus. The vertex-wise findings correspond well with reported findings for IC4, although this IC was not dominated by surface area. Lastly, linear models showed a strong and near global negative association between age and vertex-wise cortical thickness, indicating thinner cortex across almost the entire cortex with higher age. These vertex-wise findings and the differential pattern in association strength correspond well with findings reported for IC3 and also IC1

*Associations between unimodal vertex-wise structure and sex*

Permutation testing revealed moderate positive associations between sex (Supplementary Figure 8) and vertex-wise GWC, indicating that boys have higher GWC than girls in a few cortical regions including within occipital, and insular regions. These results correspond well with findings reported for IC2, although this IC was not dominated by GWC. As expected, there was a strong global positive association between sex and surface area, indicating that boys have larger surface area than girls. These vertex-wise results correspond with findings reported for IC2. Linear models showed moderate associations between sex and vertex-wise cortical thickness, indicating that boys have thinner cortex within a few dorsal brain regions such as postcentral gyrus than girls, and also thicker cortex in a few ventral brain regions such as insula. These vertex-wise results generally correspond with findings reported for IC15.

*Associations between unimodal vertex-wise structure and PGS for neuroticism*

Permutation testing revealed no significant associations between PGS for neuroticism and any of the imaging modalities.

**Supplementary Tables**

|  | Total | PING | PNC | PGS sample |
| --- | --- | --- | --- | --- |
| N | 2596 | 1129 | 1467 | 878 |
| Age (years) | 3.0-23.2  (mean= 13.8, SD= 4.6) | 3.0- 21.0  (mean= 12.1, SD= 5.1) | 8.2-23.2  (mean=15.1, SD= 3.6) | 3.17- 22.6  (mean= 14.1, SD= 4.4) |
| Sex | Boys= 1280  Girls= 1316 | Boys= 589  Girls= 540 | Boys= 691  Girls= 776 | Boys= 457  Girls= 421 |
| Self-reported ethnicity | Hispanic/Latino= 311  Native Hawaiian/Pacific Islander= 111  Asian= 274  Black/African American= 836  Native American/Alaska Native= 66  European American/White= 1476  Other=58  Not Available= 67 | Hispanic/Latino= 225  Pacific Islander= 108  Asian= 274  African American= 162  Native American= 53  White= 757  Other= Not reported  Not Available= 0 | Hispanic/Latino= 86  Native Hawaiian/Pacific Islander= 3  Asian= 0  Black/African American= 674  Native American/Alaska Native= 13  European American= 719  Other= 58  Not Available= 67 | Hispanic/Latino= 9  Native Hawaiian/Pacific Islander= 0  Asian= 1  Black/African American= 0  Native American/Alaska Native= 3  European American/White= 872  Other= 0  Not Available=0 |
| Handedness | Right handed= 2145  Left handed= 318  Mixed= 50  Not established= 11  Not available= 72 | Right handed= 948  Left handed= 118  Mixed= 50  Not established= 11  Not available= 2 | Right handed= 1197  Left handed= 200  Mixed= Not documented  Not established= Not documented  Not available= 70 | Right handed= 759  Left handed= 104  Mixed= 12  Not established= 2  Not available= 1 |

Supplementary Table 1. Sample demographics. The table shows the sample demographics for the full MRI sample, as well for PING and PNC separately. Total ethnicity numbers exceed total sample sizes as several subjects identified as belonging to more than one ethnicity group.

|  | Mean GWC (r) | Mean GWC no resid. (r) | Mean area (r) | Mean area no resid. (r) | Mean thickness (r) | Mean thickness no resid. (r) |
| --- | --- | --- | --- | --- | --- | --- |
| Scanner2 | -0.66 | -0.66 | -0.07 | -0.07 | -0.79 | -0.79 |
| Scanner7 | -0.65 | -0.65 | 0.17 | 0.17 | -0.63 | -0.63 |
| Scanner5 | -0.65 | -0.65 | 0.22 | 0.22 | -0.67 | -0.67 |
| Scanner4 | -0.67 | -0.67 | -0.06 | -0.06 | -0.78 | -0.78 |
| Scanner1 | -0.70 | -0.70 | -0.06 | -0.06 | -0.66 | -0.66 |
| Scanner3 | -0.36 | -0.36 | -0.17 | -0.17 | -0.70 | -0.70 |
| Scanner9 | -0.18 | -0.18 | -0.31 | -0.31 | -0.73 | -0.73 |
| Scanner6 | -0.70 | -0.70 | 0.30 | 0.30 | -0.64 | -0.64 |
| Scanner8 | -0.47 | -0.47 | 0.07 | 0.07 | -0.54 | -0.54 |
| Scanner10 | -0.44 | -0.44 | -0.12 | -0.12 | -0.65 | -0.65 |
| Scanner11 | -0.33 | -0.33 | -0.17 | -0.17 | -0.56 | -0.56 |

Supplementary Table 2. Pearson’s correlations of mean vertex-wise modality and age within each scanner. The table shows Pearson’s correlation coefficients of mean vertex-wise surface map values and age within each scanner, from the surface maps used in the current study as well as the same surfaces before scanner residualization (“no resid.”). Grey/white matter contrast is abbreviated as “GWC”.

| IC | Effect size (f) | P-value | IC | Effect size (f) | P-value |
| --- | --- | --- | --- | --- | --- |
| 1 | 0.43 | 1.000 | **31** | 0.05 | 1.000 |
| 2 | 0.73 | 1.000 | **32** | 0.00 | 1.000 |
| 3 | 0.10 | 1.000 | **33** | 0.01 | 1.000 |
| 4 | 4.28 | **<0.001** | **34** | 0.02 | 1.000 |
| 5 | 0.29 | 1.000 | **35** | 0.01 | 1.000 |
| 6 | 0.46 | 1.000 | **36** | 0.01 | 1.000 |
| 7 | 0.12 | 1.000 | **37** | 0.02 | 1.000 |
| 8 | 1.04 | 1.000 | **38** | 0.12 | 1.000 |
| 9 | 0.08 | 1.000 | **39** | 0.10 | 1.000 |
| 10 | 0.10 | 1.000 | **40** | 0.38 | 1.000 |
| 11 | 0.28 | 1.000 | **41** | 0.04 | 1.000 |
| 12 | 1.06 | 1.000 | **42** | 0.02 | 1.000 |
| 13 | 0.40 | 1.000 | **43** | 0.03 | 1.000 |
| 14 | 0.33 | 1.000 | **44** | 0.04 | 1.000 |
| 15 | 0.35 | 1.000 | **45** | 0.01 | 1.000 |
| 16 | 0.01 | 1.000 | **46** | 0.05 | 1.000 |
| 17 | 0.05 | 1.000 | **47** | 0.01 | 1.000 |
| 18 | 0.02 | 1.000 | **48** | 0.08 | 1.000 |
| 19 | 0.00 | 1.000 | **49** | 0.05 | 1.000 |
| 20 | 0.02 | 1.000 | **50** | 0.03 | 1.000 |
| 21 | 0.02 | 1.000 | **51** | 0.07 | 1.000 |
| 22 | 0.05 | 1.000 | **52** | 0.03 | 1.000 |
| 23 | 0.01 | 1.000 | **53** | 0.05 | 1.000 |
| 24 | 0.03 | 1.000 | **54** | 0.03 | 1.000 |
| 25 | 0.04 | 1.000 | **55** | 0.02 | 1.000 |
| 26 | 0.03 | 1.000 | **56** | 0.01 | 1.000 |
| 27 | 0.00 | 1.000 | **57** | 0.00 | 1.000 |
| 28 | 0.03 | 1.000 | **58** | 0.01 | 1.000 |
| 29 | 2.32 | 0.600 | **59** | 0.05 | 1.000 |
| 30 | 0.01 | 1.000 | **60** | 0.04 | 1.000 |

Supplementary Table 3. F-statistics of scanner effects for each independent component (IC). The table shows the F-statistic of age residualized scanner effects for each IC, as well as P-values adjusted for multiple comparisons by FDR correction. P-values at or below 0.05 are marked in bold.

**Supplementary Figures**

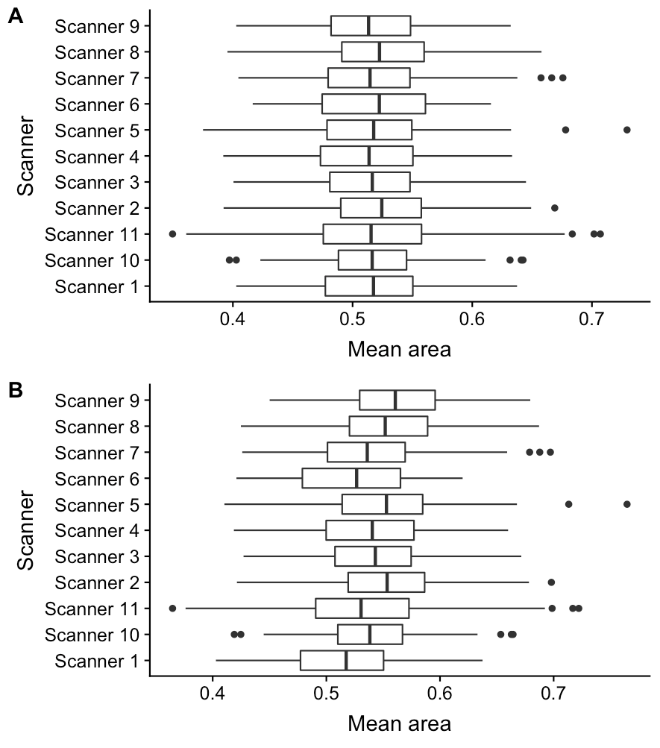

Supplementary Figure 1. Boxplots of mean vertex-wise surface area. (A) shows the mean vertex-wise surface area of the surfaces used in the current paper. (B) shows the mean vertex-wise surface area of surfaces before scanner residualization. The x- axis depicts mean area while the y-axis shows each scanner.

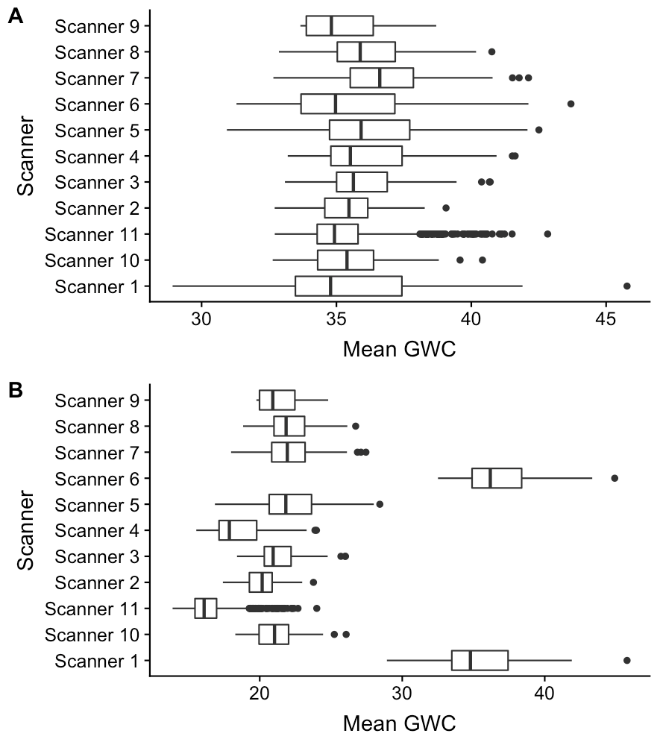

Supplementary Figure 2. Boxplots of mean vertex-wise grey/white matter contrast (GWC). (A) shows the mean vertex-wise GWC of the surfaces used in the current paper. (B) shows the mean vertex-wise GWC of surfaces before scanner residualization. The x- axis depicts mean GWC, while the y-axis shows each scanner.

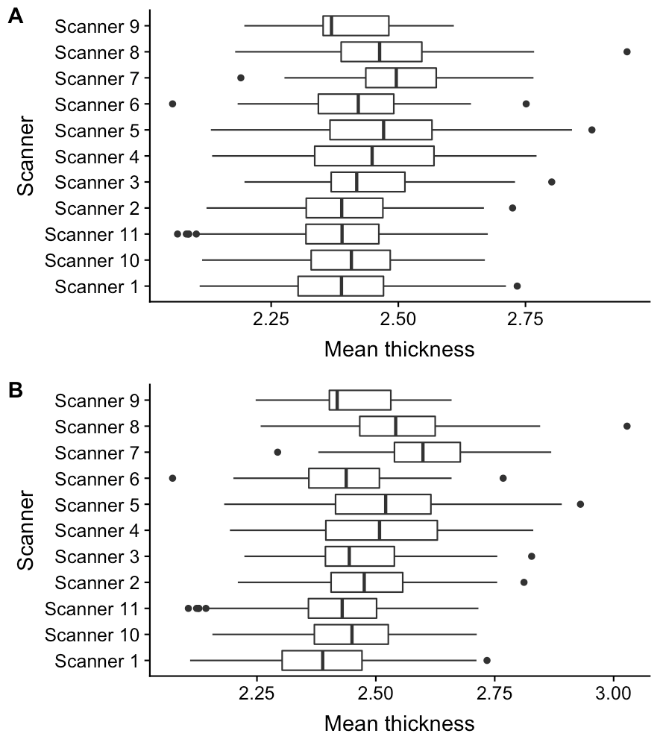

Supplementary Figure 3. Boxplots of mean vertex-wise cortical thickness. (A) shows the mean vertex-wise cortical thickness of the surfaces used in the current paper. (B) shows the mean vertex-wise cortical thickness of surfaces before scanner residualization. The x- axis depicts cortical thickness, while the y-axis shows each scanner.

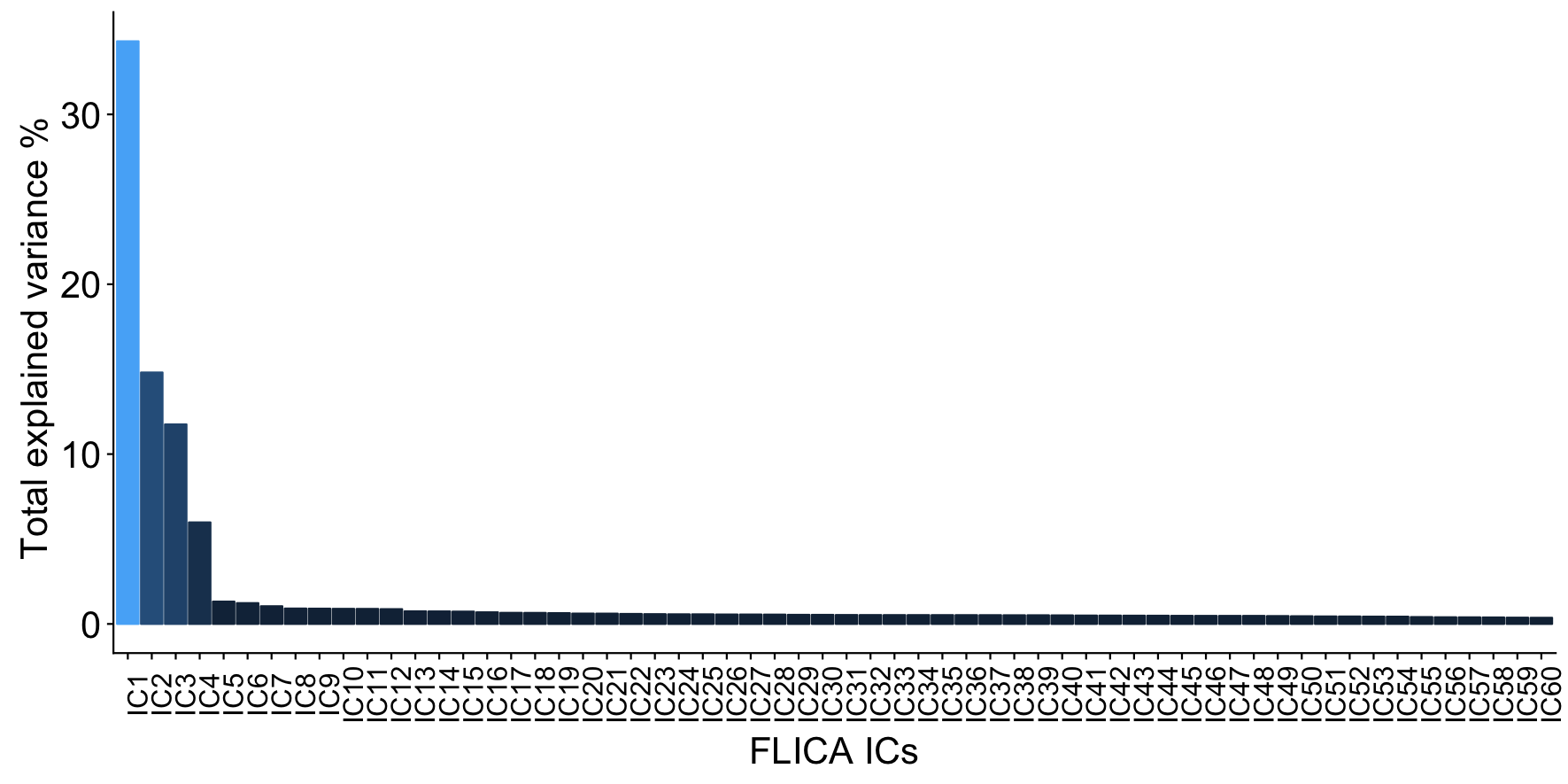

Supplementary Figure 4. Explained variance of each multimodal imaging independent component (IC). The figure shows an ordered bar plot of the percentage explained variance for each IC. The x-axis depicts each of the 60 components while the y-axis shows the total percent explained variance.

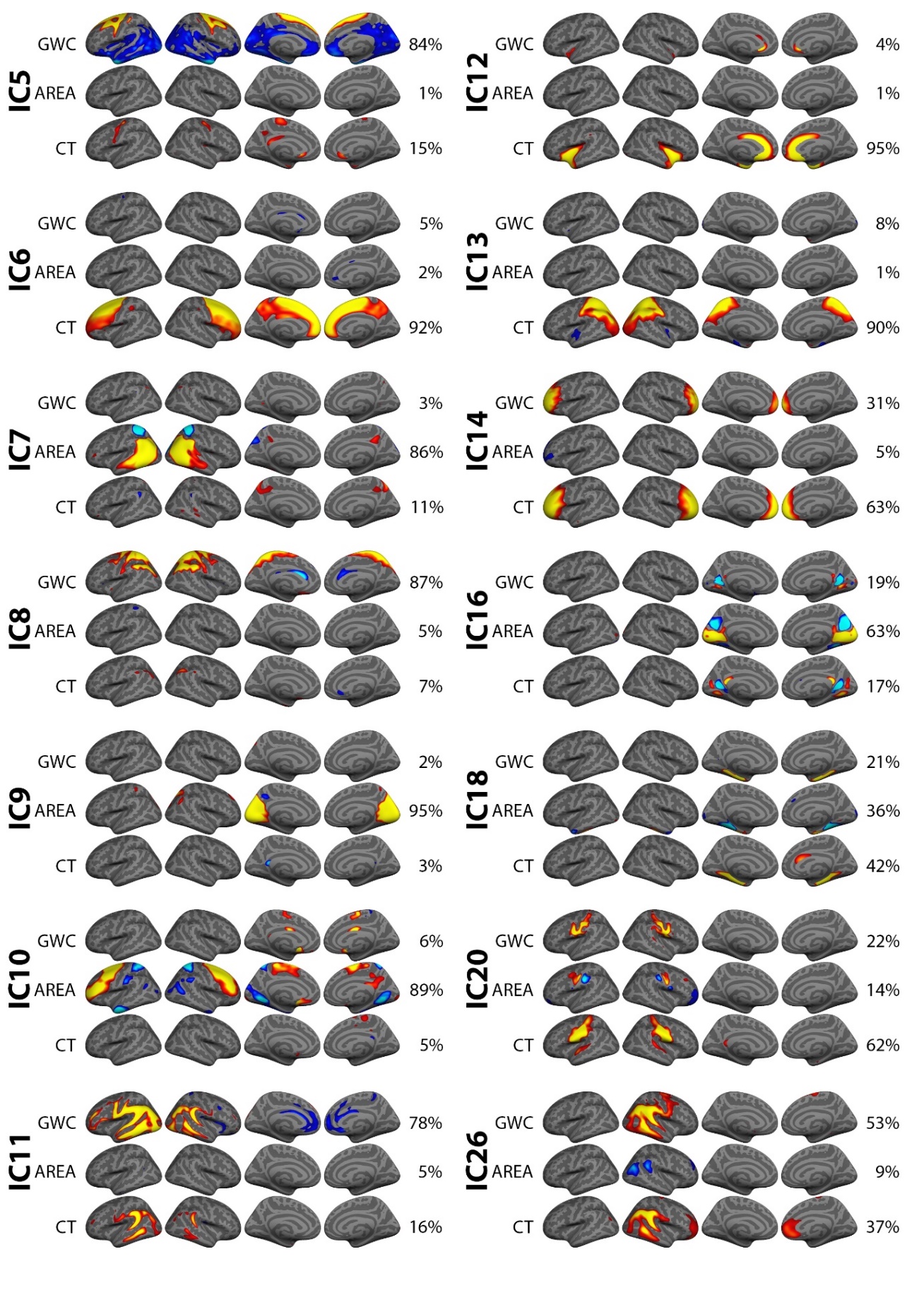

Supplementary Figure 5. Remaining independent components (ICs) showing a significant effects of age and/or sex. The figure depicts FMRIB’s linked independent component analysis (FLICA) weighted spatial maps for the first 14 of the 28 components showing a significant effects of age and/or sex, that are not depicted in the main figures of the article. All components were thresholded within a minimum and maximum of 8 and 17 standard deviations (SD) respectively.

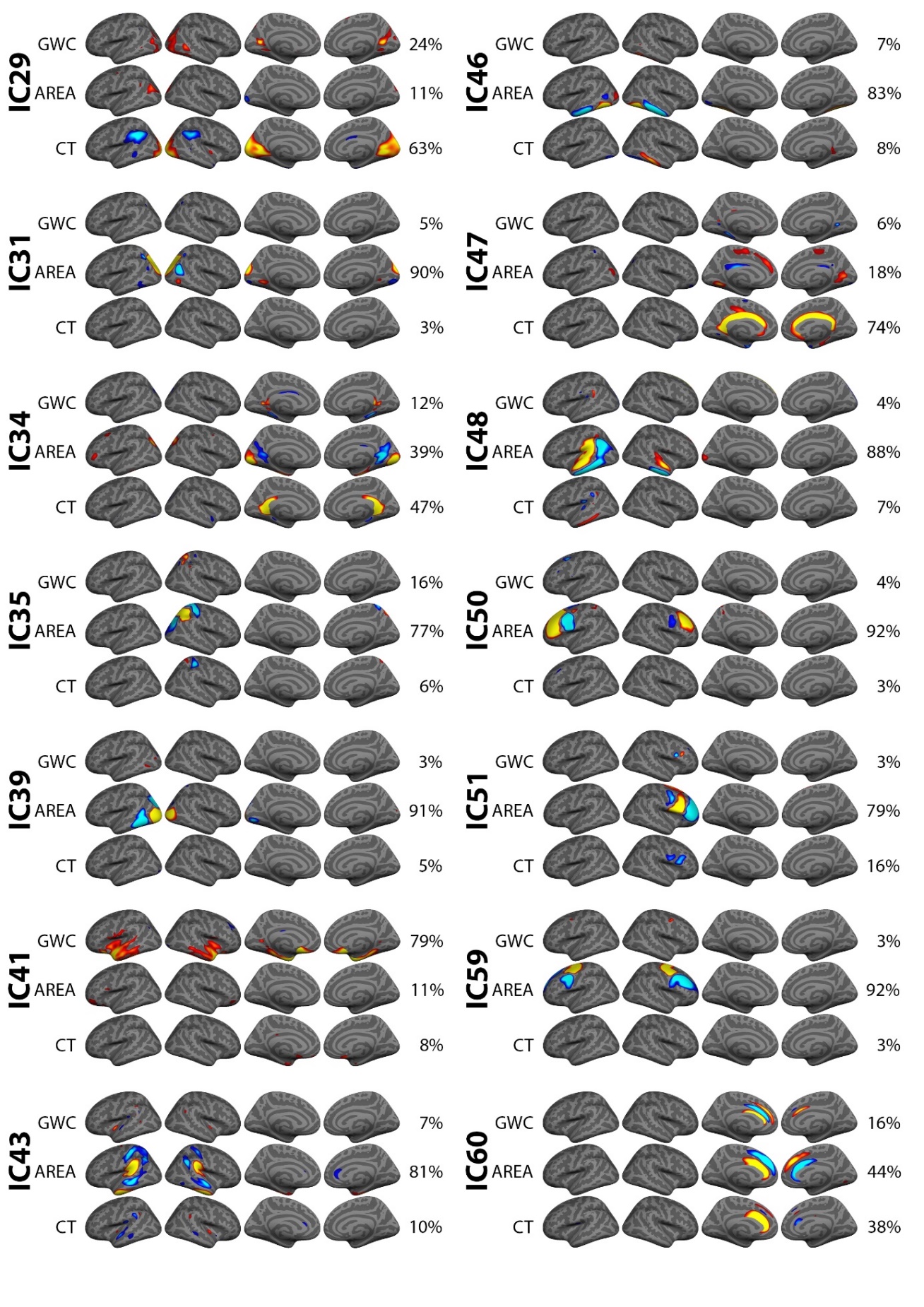

Supplementary Figure 6. Remaining independent components (ICs) showing a significant effects of age and/or sex. The figure depicts FMRIB’s linked independent component analysis (FLICA) weighted spatial maps for the latter 14 of the 28 components showing a significant effects of age and/or sex, that are not depicted in the main figures of the article. All components were thresholded within a minimum and maximum of 8 and 17 standard

deviations (SD) respectively.

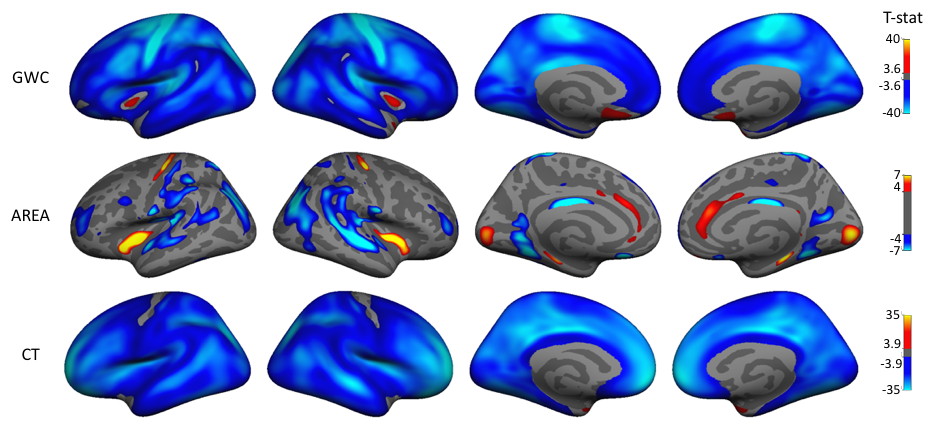

Supplementary Figure 7. Unimodal vertex-wise associations of age. The figure depicts t statistics maps, masked by familywise error corrected p values thresholded at a minimum -log_p_ of 1.6 to correct for both hemispheres. Warm colors represent a positive association and cold color represents a negative association.

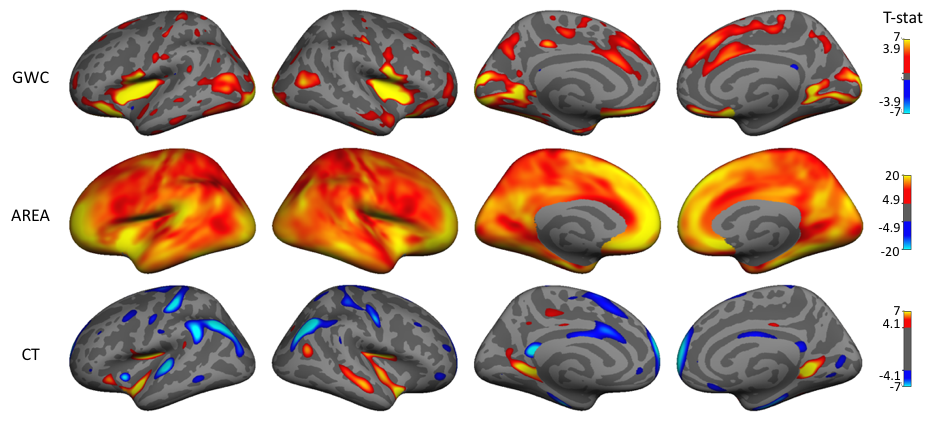

Supplementary Figure 8. Unimodal vertex-wise associations of sex. The figure depicts t statistics maps, masked by familywise error corrected p values thresholded at a minimum -log_p_ of 1.6 to correct for both hemispheres. Warm colors represent a positive association (boys > girls) and cold color represents a negative association (girls > boys).
